# Host 5-HT affects *Plasmodium* transmission in mosquitoes via modulating mosquito mitochondrial homeostasis

**DOI:** 10.1101/2024.02.19.580972

**Authors:** Li Gao, Benguang Zhang, Yuebiao Feng, Wenxu Yang, Shibo Zhang, Jingwen Wang

## Abstract

Malaria parasites hijack the metabolism of their mammalian host during the blood-stage cycle. *Anopheles* mosquitoes depend on mammalian blood to survive and to transmit malaria parasites. However, it remains understudied whether changes in host metabolism affect parasite transmission in mosquitoes. In this study, we discovered that *Plasmodium* infection significantly decreased the levels of the tryptophan metabolite, 5-hydroxytryptamine (5-HT), in both humans and mice. The reduction led to the decrease of 5-HT in mosquitoes. Oral supplementation of 5-HT to *Anopheles stephensi* enhanced its resistance to *Plasmodium berghei* infection by promoting the generation of mitochondrial reactive oxygen species. This effect was due to the accumulation of dysfunctional mitochondria caused by 5-HT-mediated inhibition of mitophagy. Elevating 5-HT levels in mouse serum significantly suppressed parasite infection in mosquitoes. In summary, our data highlight the critical role of metabolites in animal blood in determining the capacity of mosquitoes to control parasite infection.

## INTRODUCTION

Malaria, caused by infection with *Plasmodium* parasites transmitted through *Anopheles* mosquito bites, continues to be the world’s most severe parasitic disease, resulting in an estimated 247 million clinical cases and 619,000 deaths in 2021.^1^ *Plasmodium* spp. are obligate parasites that have lost multiple pathways for de novo nutrient synthesis and rely on the host for provision. Among these nutrients, amino acids are ones that parasites are auxotrophic for and largely obtain through salvage from the host.^2^ Metabolic analyses of plasma from malaria patients of different ages and disease severities reveal dysregulation in multiple amino acid metabolisms.^3^ For example, low L-citrulline and L-arginine levels have been characterized in the plasma of patients with endothelial dysfunctions.^4–6^ Elevated alanine levels are associated with lactic acidosis in severe malaria.^7^ Hyperphenylalaninemia is a well-characterized condition in both children and adults with severe and uncomplicated malaria.^3^ Tryptophan metabolism is dysregulated during *Plasmodium* infection, leading to increased levels of metabolites such as kynurenine, kynurenic acid and picolinic acid, which are positively correlated with parasitemia.^3^ Therapeutic approaches aimed at correcting the amino acid dysregulation have been shown to potentially alleviate infection pathology. For example, inhibiting the kynurenine pathway in infected mice prevents from the development of cerebral dysfunction and extends their survival.^8^ In malaria patients, arginine infusion improves endothelial function,^5^ while dietary arginine supplementation increases fetal weight and viability in an experimental mouse model of malaria in pregnancy by balancing angiogenic response and increasing placental vascularization.^9^

*Plasmodium* infection also alters the amino acid contents in mosquitoes. Mosquitoes infected with *P. berghei* exhibit increased levels of lysine, phenylalanine, proline, threonine, and tyrosine, and decreased levels of alanine, aspartic acid, glycine, and serine.^10^ Amino acid metabolism plays a crucial role in determining the susceptibility of mosquitoes to *Plasmodium* infections. The target of rapamycin (TOR) pathway that controls anabolic processes in mosquitoes by sensing the amino acid levels in the hemolymph antagonizes mosquito immune activity. Inhibition of the TOR pathway upregulates the expression of multiple immune effectors that promote parasite elimination.^11^ Prolongation of amino acids catabolism in *Anopheles* mosquitoes via silencing miR-276, which targets the branched-chain amino acid transferase, compromises the sporogony of *Plasmodium falciparum.*^12^ Additionally, the tryptophan metabolite 3-hydroxykynureine (3-HK) impairs the physical barrier, peritrophic matrix, in the midgut and facilitates *P. berghei* infection in *A. stephensi.*^13^ Another tryptophan metabolite, xanthurenic acid, acts as an exflagellation elicitor, promoting *Plasmodium*development.^14^ Therefore, *Plasmodium* infection changes the amino acid metabolism in both mammals and mosquitoes, which can impact the pathogenicity and infectivity of the parasite. However, it is currently unknown whether amino acid derangements in the host affect *Plasmodium* infection in *Anopheles* vectors during transmission from mammal to mosquito through a blood meal.

In this study, we show that 5-hydroxytryptamine (5-HT) levels are reduced in mammalian hosts (human and mice) infected with *Plasmodium* parasites. Dietary supplementation of 5-HT inhibits *P. berghei* infection in mosquitoes by promoting the generation of reactive oxygen species (ROS). The elevated ROS is a result of the accumulation of dysfunctional mitochondria due to the inhibition of mitophagy by 5-HT. We also discover that elevating 5-HT levels in mouse serum suppresses the transmission of *P. berghei* from mice to *A. stephensi*, suggesting the possibility of controlling malaria transmission by manipulating host metabolism.

## RESULTS

### The reduced 5-HT in mammalian sera facilitates *P. berghei* infection in mosquitoes

During *Plasmodium* infection in humans, the kynurenine pathway that converts tryptophan into kynurenine is perturbed.^3,15^ To get an overview of the influence of malaria parasite on tryptophan metabolism in hosts, we performed a targeted metabolomics analysis using liquid chromatography–mass spectrometry (LC– MS) (Figure 1A). Total ten malaria patients including four infected with *P. falciparum*, two with *Plasmodium vivax*, three with *Plasmodium ovale*, and one with *Plasmodium malariae*, and twelve uninfected healthy adults were included in the analysis (Table S1). Among the 15 tryptophan metabolites detected in human serum (Figure 1B), four metabolites, including L-kynurenine, quinolinic acid, cinnavalininate and 3-hydroxyl-L-kynurenine were accumulated significantly, while five metabolites, including tryptophan, 5-HT, 3-indoxyl sulfate, indole-3-propionic acid and indole acetic acid were reduced significantly in malaria patient comparing to healthy controls (Figure 1C). To investigate how *Plasmodium* infection influences tryptophan metabolism in mice, we compared tryptophan metabolism between mice 4 days post *P. berghei* infection and age-matched non-infected controls. Out of the 10 metabolites detected, four showed significant alterations. Among these metabolites, the serum levels of L-kynurenine and cinnabarinic acid were significantly elevated, while the levels of 5-HT and 5-hydroxytryptophan were decreased in *P. berghei* infected mice (Figures 1D and 1E). Since 5-HT was decreased significantly in both human and mice infected with different species of *Plasmodium*, we speculated that *Plasmodium* infection might similarly reduce 5-HT levels in mosquitoes. As expected, the 5-HT level was significantly reduced in the midguts of mosquitoes that were supplied with a blood meal containing *P. berghei*, compared to the ones ingested un-infectious blood (Figure 1F). However, when the blood bolus was removed from mosquito midgut, the 5-HT levels remained comparable between the two groups (Figure S1A). We next examined the 5-HT levels in mosquitoes 3-day post blood meal when the blood was digested completely and found a significant reduction of 5-HT in mosquitoes infected with parasites (Figure S1B). These results indicate that the mosquito 5-HT levels were determined by dietary 5-HT. Altogether, these results suggest that *Plasmodium* infection significantly reduces 5-HT levels in mammalian hosts (human and mice), and this disturbance could be transmitted to mosquito vectors (*A. stephensi*) through a blood meal.

**Figure 1.**
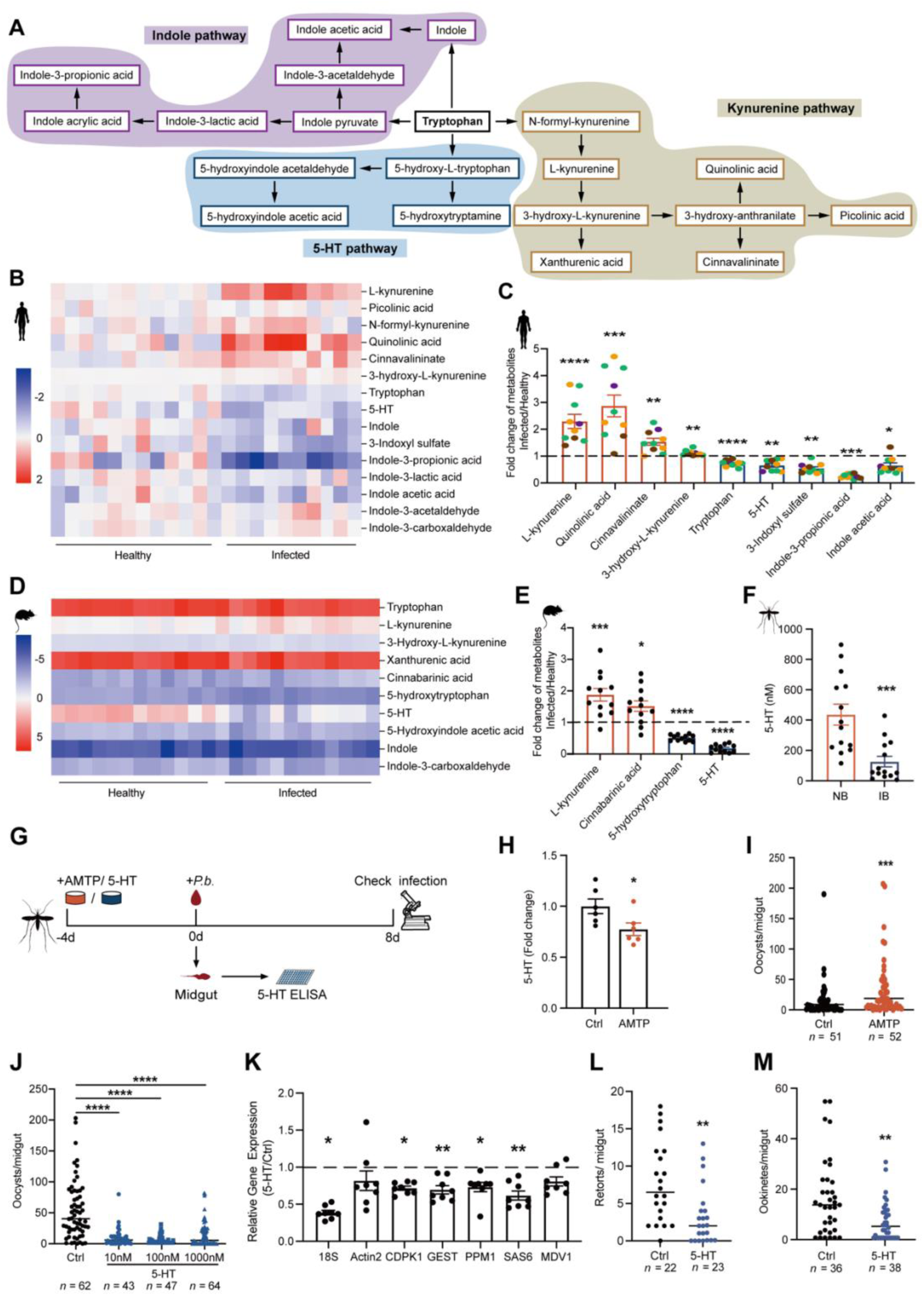
5-HT inhibits *P. berghei* infection in mosquitoes. (A) Overview of tryptophan metabolic pathway in mammalian hosts. The purple boxes represent the metabolites of the indole pathway, the blue boxes represent the metabolites of the 5-HT pathway, and the tan boxes represent the metabolites of the kynurenine pathway. (B) Heatmap of 15 tryptophan and metabolites detected in the sera of healthy (Healthy) and parasite infected adults (Infected). Increased metabolites were shown in red; decreased metabolites were shown in blue. (C) Fold changes of differentially altered metabolites in the sera of malaria patients (Infected, *n* = 10) versus healthy adults (Healthy, *n* = 12) analyzed by LC–MS. Metabolite abundance in malaria patients was normalized to that in healthy adults. Each dot represented an individual adult infected with *P. ovale* (orange), *P. falciparum* (green), *P. vivax* (brown) or *P. malariae* (purple). Data were shown as mean ± SEM. (D) Heatmap of 10 tryptophan-metabolites detected in the sera of non-infected (Healthy) and *P. berghei* infected (Infected) mice. Increased metabolites were shown in red; decreased metabolites were shown in blue. (E) Fold changes of differentially altered metabolites in the sera of *P. berghei* infected (Infected, *n* = 12) versus non-infected (Healthy, *n* = 12) mice analyzed by LC–MS. Metabolite abundance in *P. berghei* infected mice was normalized to that of non-infected ones. Each dot represented an individual mouse. Data were shown as mean ± SEM. (F) The concentrations of 5-HT in the mosquito midguts 24 h post normal blood (NB, *n* = 14) and *P. berghei* infected blood (IB, *n* = 14) analyzed by ELISA. Thirty midguts were pooled for one biological sample. Each dot represented one biological replicate. Data were pooled from three independent experiments and shown as mean ± SEM. (G) Schematic overview of AMTP and 5-HT supplementation in mosquitoes. (H) Fold change of 5-HT in mosquitoes treated with (AMTP, *n* = 6) and without (Ctrl, *n* = 6) 100 μM AMTP for 4 days (prior to blood feeding) analyzed by ELISA. The abundance of 5-HT in AMTP treated mosquitoes was normalized to that in controls. Twenty-five mosquitoes were pooled for one biological sample. Each dot represented one biological replicate. Data were pooled from two independent experiments and shown as mean ± SEM. (I) Oocyst numbers in control (Ctrl, *n* = 51) and AMTP (AMTP, *n* = 52) treated mosquitoes. Each dot represented an individual mosquito. Data were pooled from two independent experiments and horizontal lines represented the medians. (J) Oocyst numbers in control (Ctrl, *n* = 62) and mosquitoes orally supplemented with 10 nM (*n* = 43), 100 nM (*n* = 47) and 1000 nM (*n* = 64) 5-HT. Each dot represented an individual mosquito. Data were pooled from two independent experiments and horizontal lines represented the medians. (K) Fold changes of male gametogenesis associated genes in the midguts of control (*n* = 8) and 1 μM 5-HT treated (*n* = 8) mosquitoes 15 min post infection. The expression levels of the target genes were normalized to *S7*. The relative expression levels of target genes in mosquitoes treated with 5-HT were normalized to those in the control group. Results from one of two independent experiments were shown. The second replication was shown in Fig S1G. Each dot represented five mosquito midguts. Data were shown as mean ± SEM. (L) Retort numbers in the midguts of control (*n* = 36) and 1 μM 5-HT (*n* = 38) treated mosquitoes 12 h post infection. Each dot represented an individual mosquito. Data were pooled from two independent experiments and horizontal lines represented the medians. (M) Ookinete numbers in the midguts of control (*n* = 36) and 1 μM 5-HT (*n* = 38) treated mosquitoes 24 h post infection. Each dot represented an individual mosquito. Data were pooled from two independent experiments and horizontal lines represented the medians. Significance was determined by two-sided Student’s t test in (C), (E), (F), (H) and (K), Mann-Whitney test in (I), (L) and (M) and ANOVA with Dunn’s test in (J). *p < 0.05, **p < 0.01, ***p < 0.001, ****p < 0.0001.

To investigate whether 5-HT plays a role in influencing *P. berghei* infection in mosquitoes, we orally administrated mosquitoes with a-methyl-DL-tryptophan (AMTP), an antagonist of tryptophan hydroxylase (TPH), which is the rate-limiting enzyme of 5-HT biosynthesis, for four days prior to *P. berghei* infection (Figure 1G). By blocking TPH, we inhibited the biosynthesis of 5-HT (Figure 1H), and observed a significant increase in oocyst number in mosquitoes (Figure 1I). We next raised mosquito 5-HT levels by orally supplementing increased amounts of 5-HT with sugar meal for four days prior to blood feeding (Figures 1G and S1F). We determined the amount of 5-HT to use based on its levels in mosquitoes, as well as in the blood of humans (Figures 1F and S1C). All three doses led to a significant decrease in oocyst numbers (Figure 1J). Since 1 µM is a similar concentration to that found in healthy human blood (Figure S1C),^16^ and oral supplementation of 1 µM 5-HT to mosquitoes didn’t affect the amount of blood mosquito intake (Figures S1D and S1E), we used this concentration for the following treatment. After being ingested by mosquitoes, *Plasmodium* forms gametes within about 15 minutes. The gametes then undergo fertilization and differentiate into retorts, ookinetes, and oocysts^17^. We then examined which developmental stage was impaired by 5-HT by quantifying male gametogenesis via qPCR 15 min post-infection,^18^ and counting retort and ookinete numbers microscopically 12 h and 24 h post-infection respectively. The administration of 5-HT significantly reduced the numbers of gametes, retorts and ookinetes compared with controls (Figures 1K-1M), suggesting that 5-HT exerts a killing effect on *P. berghei* right after parasites’ arrival in midgut. Altogether, these results indicate that a decrease in 5-HT levels in the host’s serum might promote the transmission of *Plasmodium* in mosquitoes.

### Mitochondrial ROS inhibits *Plasmodium* infection

Peripheral 5-HT plays an important role in regulating the immune system in the mammalian gut.^19,20^ To examine whether 5-HT affects mosquito immune responses, we measured the expression levels of major immune genes and genes related to reduction-oxidation (redox) reactions in midguts 24 h post infection. The mRNA levels of most immune genes were not significantly altered by 5-HT administration. However, the expression of three genes encoding antioxidant enzymes, including copper-zinc superoxide dismutase 3 (CuSOD_3_), catalase 1 (CAT), and uridine 5’-diphospho-glycoprotein glucosyltransferase (UGT) were significantly upregulated, indicating the change in the redox state of the midgut (Figure 2A). We next examined the reactive oxygen species (ROS) levels of midguts microscopically by staining with dihydroethidium (DHE), a superoxide indicator. As expected, 5-HT supplementation significantly increased superoxide levels in mosquitoes 24 h and 15 min post infection (Figures 2B and S2A). ROS is a potent anti-*Plasmodium* agent in *Anopheles* mosquitoes.^21^ To assess whether 5-HT-induced ROS inhibits parasite infection, we scavenged ROS by simultaneously supplementing 5-HT and vitamin C to mosquitoes. Vitamin C effectively reduced midgut ROS (Figure 2B) and restored oocyst numbers to control level (Figure 2C). Similarly, H_2_O_2_ level was increased significantly in 5-HT-treated mosquitoes and restored to control level when vitamin C was added (Figure S2B). Administration of H_2_O_2_ had the same inhibitory effect in *P. berghei* infection as 5-HT did (Figures S2C-S2G).

**Figure 2.**
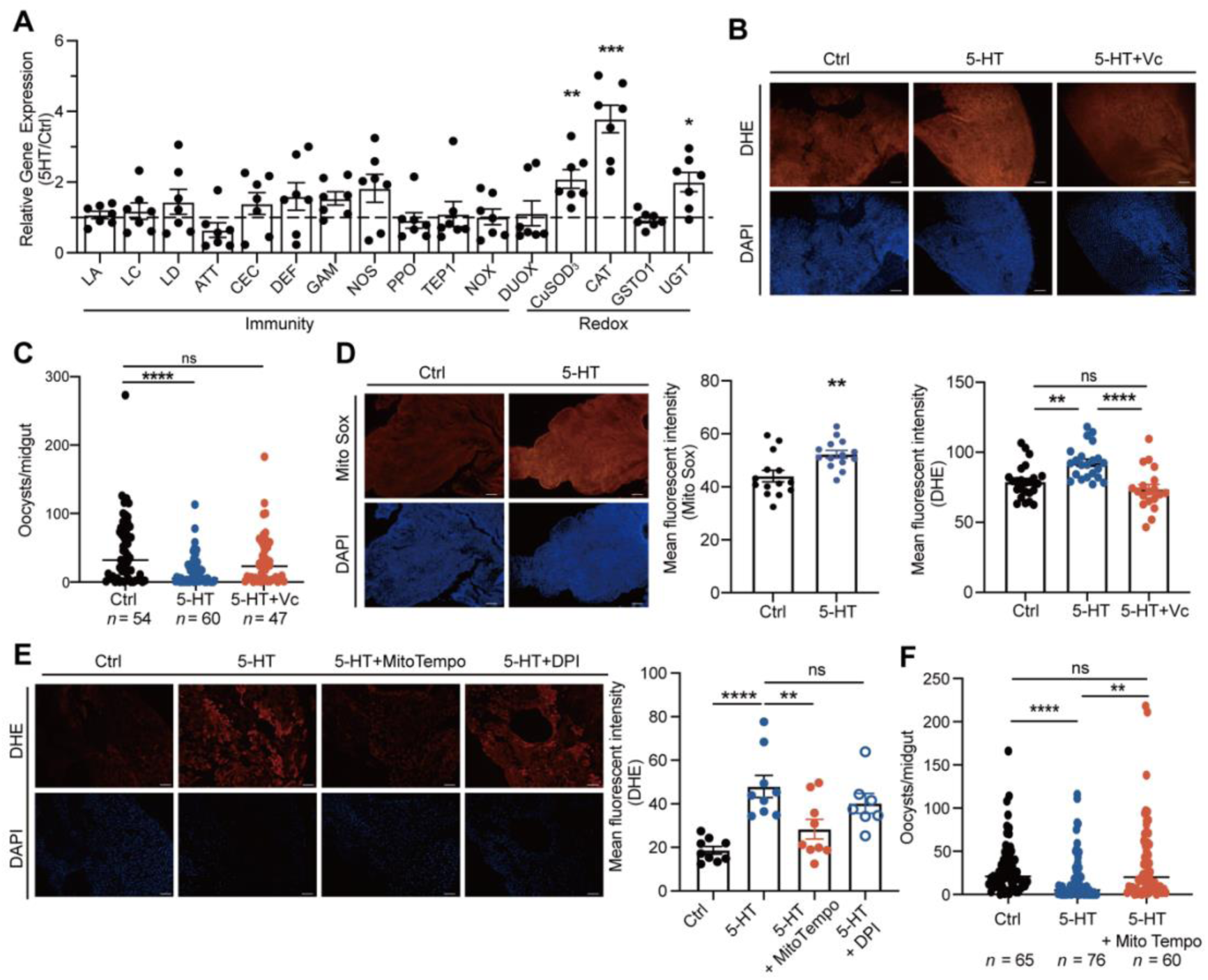
5-HT- induced mitochondrial ROS inhibits *P. berghei* infection. (A) Fold changes of immune- and redox-related genes in the midguts of control (*n* = 7) and 1 μM 5-HT treated (*n* = 7) mosquitoes 24 h post infection. The expression level of the target gene was normalized to *S7*. The relative expression levels of target genes in 5-HT treated mosquitoes were normalized to those in controls. Each dot represented an individual mosquito midgut. Results from one of two independent experiments were shown and data were shown as mean ± SEM. The second replication was shown in Figure S2. (B) DHE staining (red) in the midguts of control, 5-HT (1 μM) and 5-HT+vitamin C (3.3 mM) treated mosquitoes 24 h post infection. Nuclei were stained with DAPI (blue). Representative images were shown (up). Mean fluorescent intensity was measured and calculated as described (low). Each dot represented an individual mosquito midgut. Data were pooled from three independent experiments and shown as mean ± SEM. Scale bar, 25 μm. (C) Oocyst numbers in the midguts of mosquitoes supplemented with 5-HT (*n* = 60), 5-HT and vitamin C simultaneously (5-HT+Vc, *n* = 47) and controls (Ctrl, *n* = 54). Each dot represented an individual mosquito. Data were pooled from two independent experiments and horizontal lines represented the medians. (D) MitoSOX (red) staining in the midgut of control and 5-HT treated mosquitoes 24 h post infection. Nuclei were stained with DAPI (blue). Representative images were shown (left). Mean fluorescent intensity was measured and calculated (right). Each dot represented an individual mosquito midgut. Data were pooled from two independent experiments and shown as mean ± SEM. Scale bar, 25 μm. (E) DHE (red) staining in the midgut of mosquitoes treated with 5-HT, 5-HT+ MitoTempo (50 μM) and 5-HT+ DPI (50 μM) and control midguts 24 h post infection. Nuclei were stained with DAPI (blue). Representative images were shown (left). Mean fluorescent intensity was measured and calculated (right). Each dot represented an individual mosquito midgut. Data were pooled from two independent experiments and shown as mean ± SEM. Scale bar, 25 μm. (F) Oocyst numbers in the midguts of control (*n* = 65), 5-HT (*n* = 76) and 5-HT + MitoTempo (*n* = 60) treated mosquitoes. Each dot represented an individual mosquito. Data were pooled from two independent experiments and horizontal lines represented the medians. Significance was determined by two-sided Student’s t test in (A) and (D) and ANOVA with Dunn’s test in (C) and Tukey’s test in (B), (E) and (F). *p < 0.05, ***p < 0.001, ****p < 0.0001, ns, not significant.

ROS is generated by NADPH oxidases including nicotinamide adenine dinucleotide phosphate oxidase (NOX) and Dual oxidases (DUOX), as well as a byproduct of mitochondrial oxidative phosphorylation.^22,23^ As expression levels of *NOX* and *DUOX* showed no significant difference between 5-HT treated and control mosquitoes (Figure 2A), we hypothesized that mitochondria might be responsible for the increased ROS generation. We next measured mitochondrial-derived ROS in midguts using MitoSOX red staining and found increased fluorescent signals in 5-HT supplemented mosquitoes 24 h and 15 min post infection (Figure 2D and S2H). To investigate whether 5-HT specifically promotes mitochondrial ROS generation, we inhibited mitochondrial- and NOX-derived ROS by MitoTEMPO and DPI (dibenziodolium chloride), respectively.^24^ As expected, ROS was scavenged in mosquitoes treated with MitoTEMPO but not with DPI (Figure 2E). Accordingly, the addition of MitoTEMPO, but not DPI restored oocyst numbers to control levels (Figures 2F and S2I). Therefore, these results suggest that oral administration of 5-HT promotes the production of mitochondrial ROS, leading to the elimination of *P. berghei* in mosquitoes.

### Accumulation of damage mitochondria increases ROS production

The elevation of mitochondrial ROS is associated with the increased mitochondrial biogenesis or damage. To determine whether dietary 5-HT promotes mitochondrial biogenesis, we quantified the amount of mitochondrial DNA (MtDNA) and proteins by using the five mitochondrial-encoded genes, including cytochrome c oxidase subunit I and II (*COX1* and *COX2*) and NADH dehydrogenase 1, 2 and 4 (*ND1*, *2*, and *4*), and two proteins, including a mitochondrial inner membrane protein ATP synthase F1 subunit alpha (ATP5A) and an outer membrane protein TOMM20 as indicators, respectively. We did not observe significant differences in the levels of MtDNA or proteins between 5-HT-treated midguts and non-treated controls (Figures 3A-3C), suggesting that 5-HT-induced ROS production is not a result of increased mitochondrial biogenesis.

**Figure 3.**
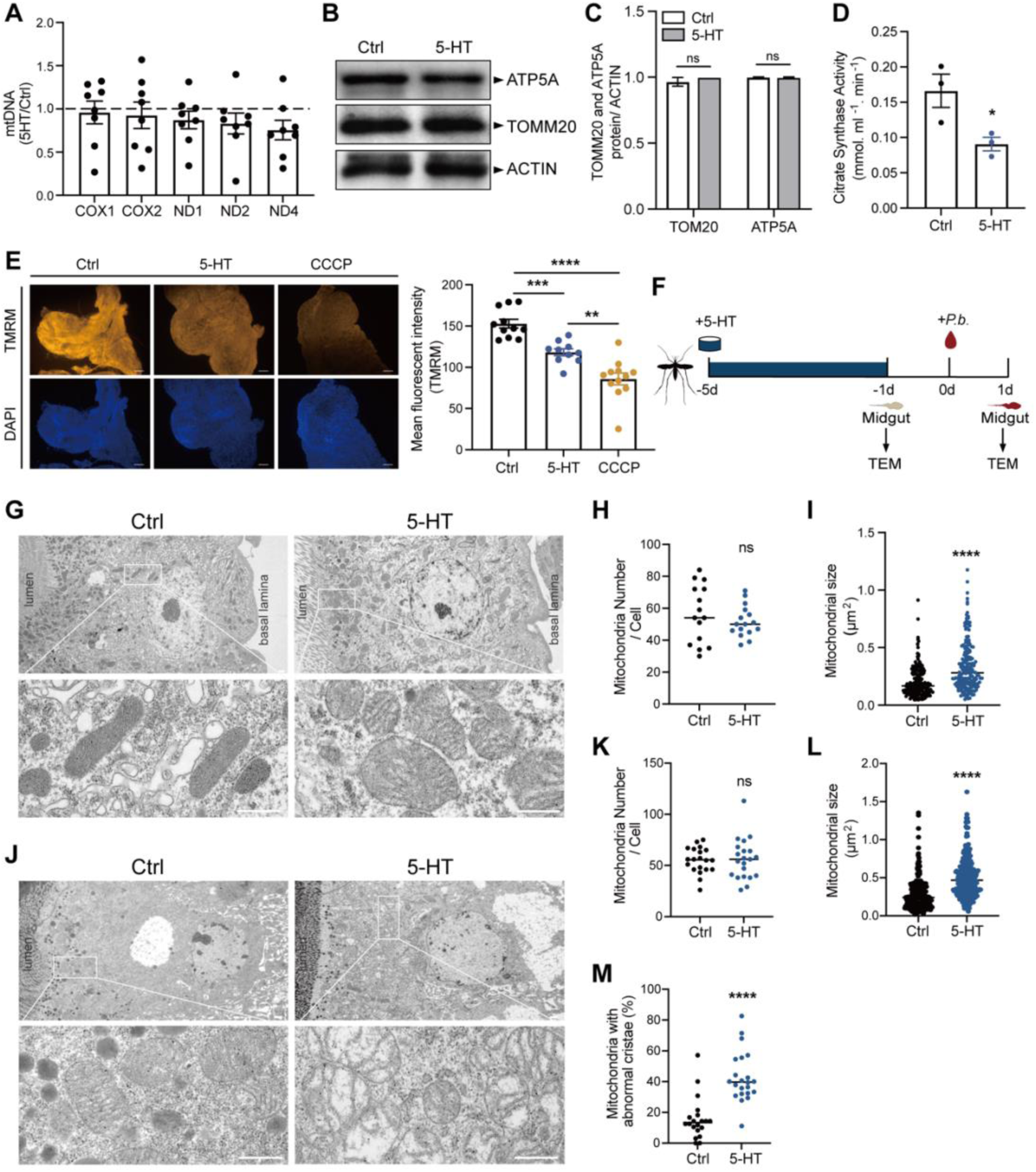
5-HT supplementation induces mitochondrial damage. (A) Relative mtDNA expression levels in the midgut of control (*n* = 8) and 5-HT treated (*n* = 8) mosquitoes 24 h post infection. The expression level of the target gene was normalized to *S7*. The relative mtDNA expression levels in 5-HT treated mosquitoes were normalized to that in controls. Each dot represented an individual mosquito midgut. Results from one of two independent experiments were shown. The second replication was shown in Figure S3A. Data were shown as mean ± SEM. (B) Western blot of TOMM20 and ATP5A in the midgut of control and 5-HT treated mosquitoes 24 h post infection. (C) Quantification of band intensities in (B). The expression level of the target protein was normalized to ACTIN. Data were pooled from three independent experiments and shown as mean ± SEM. (D) Citrate synthesis activity in the midguts of control (*n* = 3) and 5-HT treated (*n* = 3) mosquitoes 4 days post treatment. A hundred mosquito midguts were pooled for one sample. Each dot represented an individual biological replicate. Data were pooled from three independent experiments and shown as mean ± SEM. (E) The mitochondrial membrane potential measured via TMRM (red) staining in the midgut of control and 5-HT treated mosquitoes 24 h post infection. The mitophagy inducer, carbonyl cyanide m-chlorophenyl hydrazone (CCCP), was used as a positive control. The nuclei were stained with DAPI (blue). Representative images were shown (left). Mean fluorescence intensity was measured and calculated (right). Each dot represented an individual mosquito midgut. Data were pooled from two independent experiments and shown as mean ± SEM. Scale bar, 25 μm. (F) Schematic overview of TEM at two time points following 5-HT supplementation in mosquitoes. (G-I) Mitochondrial morphology in control and 5-HT supplemented mosquitoes 4 days post 5-HT treatment. Mitochondrial structure, higher magnification images of the white boxed regions are shown in the lower panels (G), number (H) and size (I) were evaluated by transmission electron micrographs (TEM). Each dot represented an individual midgut cell in (H), individual mitochondria in (I). Horizontal lines represented the medians. Images are representatives of 15 midguts per group. Scale bar, 500 nm. (J-M) Mitochondrial morphology in control and 5-HT supplemented mosquitoes 24 h post infection. Mitochondrial structure, higher magnification images of the white boxed regions are shown in the lower panels (J), number (K), size (L) and the ratio of mitochondria with abnormal cristae (M) were evaluated by transmission electron micrographs (TEM). Each dot represented an individual midgut cell in (K) and (M), individual mitochondria in (L). Horizontal lines represented the medians. Images are representatives of 20 midguts per group. Scale bar, 500 nm. Significance was determined by two-sided Student’s t test in (A), (C), (D) and (E) and Mann-Whitney test in (H), (I) and (K-M). *p < 0.05, **p < 0.01, ***p < 0.001, ****p < 0.0001, ns, not significant.

We next investigated whether the 5-HT supplementation affects mitochondrial function by analyzing the enzymatic activity of citrate synthase, a key enzyme in the Krebs cycle. We observed a 45.8% reduction in citrate synthase activity in the midguts of mosquitoes treated with 5-HT (Figure 3D). Mitochondrial membrane potential (ΔΨm) is another indicator of mitochondrial activity. We found that mitochondrial membrane potential in mosquitoes treated with 5-HT was moderately but significantly reduced, as measured by the probe, methyl ester (TMRM) (Figure 3E). Similar results were observed in 5-HT treated-MSQ43 cells derived from *A. stephensi* (Figure S3A). We also monitored mitochondrial respiration by measuring their oxygen consumption rate (OCR).^25^ However, due to the difficulties in collecting enough mitochondria from mosquito midguts, we switched to MSQ43 cells for OCR analysis. As expected, the addition of 5-HT to MSQ43 cells reduced the basal respiration (by 36.8%), ATP-linked respiration (by 37.3%), maximal respiration (by 32.6%) and extra respiration (by 17.6%) (Figure S3B).

To further confirm the influence of 5-HT treatment on mitochondrial dysfunction, we examined mitochondrial morphology at two time points, 5-HT treatment for 4 days (24 h prior to *P. berghei* infection) and 24 h post *P. berghei* infection, using transmission electron microscopy (Figure 3F). In both blood-unfed and - fed midguts, mitochondria accumulated in the apical site of epithelial cells close to the midgut lumen (Figures 3G and 3J). The administration of 5-HT didn’t affect the number of mitochondria at either time point (Figures 3H and 3K). However, it did increase mitochondrial size by 67.3% 24 h prior to and by 82% 24 h post infection, compared to controls (Figures 3I and 3L). It is noteworthy that the well-organized stacks of cristae typically found in healthy mitochondria were replaced by sparse and fragmented cristae following the administration of 5-HT 24 h post infection (Figure 3J). Furthermore, some mitochondria exhibited large and vacant central matrix spaces (Figure 3J). The proportion of mitochondria with abnormal cristae rose by 27.1 % in 5-HT supplemented mosquitoes compared to controls (Figure 3M). These findings collectively suggest that 5-HT supplementation impairs mitochondrial function in mosquito midguts, leading to heightened ROS production.

### 5-HT inhibits mitophagy

The accumulation of functional compromised mitochondria in 5-HT treated mosquitoes might be due to the failure to eliminate unhealthy mitochondria.^26^ Mitophagy is a mitochondrial quality control mechanism, in which dysfunctional mitochondria are engulfed by autophagosomes and fused with lysosomes for degradation.^27^ To test whether 5-HT inhibits mitophagy, we first accessed mitophagy in MSQ43 cells treated with 5-HT by co-staining the outer mitochondrial membrane protein TOMM20 and the autophagic microtubule-associated protein 1 light chain 3B (LC3B), a member of the ATG8 family that are involved in autophagosome development and maturation.^28^ Cells treated with CCCP, an inducer of mitophagy were used as a positive control. Treatment with 5-HT significantly reduced the colocalization of mitochondria with autophagosomes compared to control group (Figure 4A). Additionally, it decreased the formation of LC3 puncta (Figure 4B) and the association of LC3 puncta with mitochondria (Figure 4C), indicating that 5-HT inhibits mitophagy in vitro. We next tested the effect of 5-HT on lysosome-mitochondria association in vivo by staining freshly dissected midguts with Mitotracker and Lysotracker, and observed the similarly inhibitory effects of 5-HT on mitophagy (Figure 4D). Consistently, the protein level of LC3 was reduced upon the addition of 5-HT (Figures 4E and 4F). Altogether, these results indicate that oral administration of 5-HT inhibits mitophagy in midguts.

**Figure 4.**
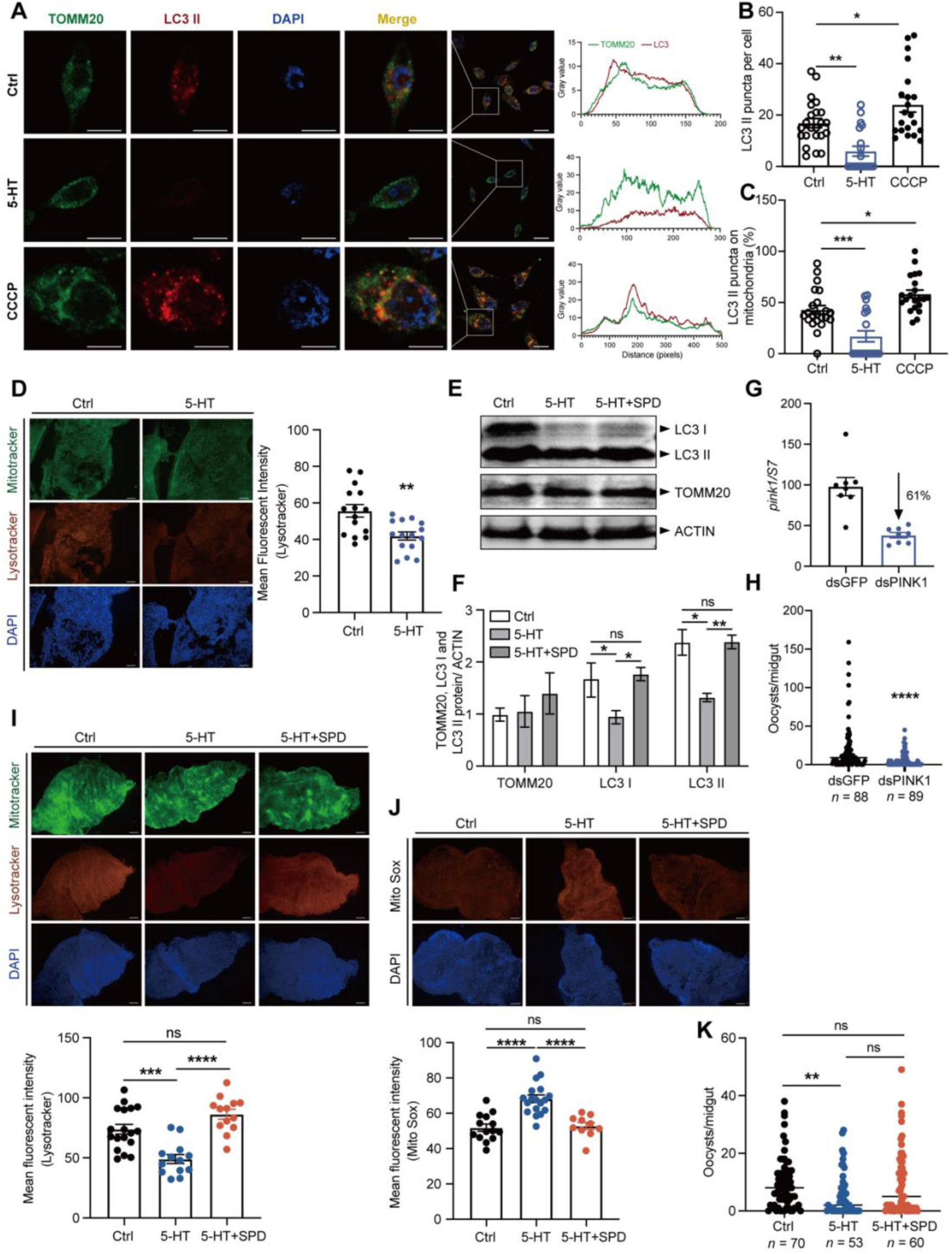
5-HT supplementation inhibits mitophagy. (A) Immunostaining of mitochondria (TOMM20, green) and autophagosome (LC3 II, red) in control and 5-HT treated MSQ43 cells 4 days post treatment. Nuclei were stained with DAPI (blue). Cells treated with 50 nM CCCP for 20 min were used as a positive control. The Pearson’s coefficient indexes between LC3 II-red and TOMM20-green fluorescence intensities were determined in 10 or more cells from three independent experiments. Scale bar, 5 μm. (B) Calculation of LC3 puncta in control (*n* = 23), 5-HT (*n* = 19) and CCCP (*n* = 21) treated MSQ43 cells in A. Each dot represented an individual cell. Data were pooled from three independent experiments and shown as mean ± SEM. (C) The ratio of LC3 puncta colocalized with mitochondria in control (*n* = 23), 5-HT (*n* = 19) and CCCP (*n* = 21) treated MSQ43 cells in (A). Each dot represented an individual cell. Data were pooled from three independent experiments and shown as mean ± SEM. (D) Co-staining of Mitotracker (green) and Lysotracker (red) in the midgut of control and 5-HT treated mosquitoes 24 h post infection. Nuclei were stained with DAPI (blue). Representative images were shown (left). Mean fluorescence intensity of Lysotracker was measured and calculated as described in Methods (right). Each dot represented an individual mosquito midgut. Data were pooled from three independent experiments and shown as mean ± SEM. Scale bar, 25 μm. (E) Western blot of LC3 I, LC3 II and TOMM20 in the control, 5-HT and 5-HT + spermidine (SPD, 100 μM) treated mosquitoes 24 h post infection. (F) The quantification of band intensities in (E). The expression level of the target protein was normalized to ACTIN. Data were pooled from three independent experiments and shown as mean ± SEM. (G) The *pink1* silencing efficiency in mosquitoes. Expression level of *pink1* was normalized to *A*. *stephensi S7*. Relative expression level of *pink1* in dsPINK1 mosquitoes was normalized to that in dsGFP controls. Each dot represented an individual mosquito. The data were shown as mean ± SEM. (H) Oocyst numbers in the midguts of dsGFP (*n* = 88) and dsPINK1(*n* = 89) mosquitoes. Each dot represented an individual mosquito. Data were pooled from three independent experiments and horizontal lines represented the medians. (I) Co-staining of Mitotracker (green) and Lysotracker (red) in the midgut of control, 5-HT and 5-HT + SPD treated mosquitoes 24 h post infection. Nuclei were stained with DAPI (blue). Representative images were shown (left). Mean fluorescence intensity of Lysotracker was measured and calculated (right). Each dot represented an individual mosquito midgut. Data were pooled from two independent experiments and shown as mean ± SEM. Scale bar, 25 μm. (J) Mito-Sox (red) staining in the midgut of control, 5-HT and 5-HT + SPD treated mosquitoes 24 h post infection. Nuclei were stained with DAPI (blue). Representative images were shown (left). Mean fluorescence intensity was measured and calculated (right). Each dot represented an individual mosquito midgut. Data were pooled from two independent experiments and shown as mean ± SEM. Scale bar, 25 μm. (K) Oocyst numbers in the midguts of control (*n* = 70), 5-HT (*n* = 53) and 5-HT + SPD (*n* = 60) treated mosquitoes. Each dot represented an individual mosquito. Data were pooled from two independent biological experiments and horizontal lines represented the medians. Significance was determined by ANOVA with Dunnett’s test in (B), (C), Tukey’s test in (H) and (I) and Dunn’s test in (J), two-sided Student’s t test in (D) and (F) and Mann-Whitney test in (G). *p < 0.05, **p < 0.01, ***p < 0.001, ****p < 0.0001, ns, not significant.

We next examined whether inhibition of mitophagy replicates the effects of 5-HT on parasite infection. PINK1 (Phosphatase and tensin homologue (PTEN) - induced kinase 1) is responsible for the initiation of mitophagy.^27^ We then inhibited mitophagy via knocking down *PINK1*. As expected, knockdown of *PINK1* significantly induced ROS generation (Figures 4G and S4A) and inhibited parasite infection (Figures 4H and S4B). We next rescued 5-HT-mediated mitophagy by simultaneously administrating spermidine, an activator of mitophagy, and 5-HT to mosquitoes.^29,30^ Addition of spermidine restored the levels of LC3 protein, mitochondrial ROS, and mitophagy activity (Figures 4E, 4F, 4I, and 4J). It also moderately increased susceptibility of mosquitoes to parasite infection but without statistical significance (Figure 4K). One possible explanation is that the concentration and timing of spermidine supplementation may not have been optimal for inducing sustained mitophagy in the mosquito midgut. Taken together, our results show that 5-HT-mediated inhibition of mitophagy leads to the increased dysfunctional mitochondria. This disruption of mitochondrial homeostasis in the mosquito midgut ultimately affects the infection outcomes of *Plasmodium*.

### Elevating 5-HT in mice serum inhibits *Plasmodium* infection in mosquitoes

Given that increasing 5-HT intake through a sugar meal inhibits *P. berghei* infection in mosquitoes, we evaluated the possibility of manipulating 5-HT levels in *P. berghei*-infected mice to control parasite transmission in mosquitoes. We first orally supplemented the 5-HT reuptake inhibitor fluoxetine to mice through drinking water at the same day when they were infected with *P. berghei*, with saline solution used as negative controls (Figure 5A). After four days, serum 5-HT levels were measured, and mosquitoes were allowed to feed on these mice. Oral administration of fluoxetine increased serum 5-HT levels compared to controls (Figure 5B) and accordingly significantly inhibited *P. berghei* infection in mosquito midguts (Figure 5C). We next examined whether continuously feeding fluoxetine to mice would alleviate *Plasmodium* pathogenicity. Unexpectedly, administration of fluoxetine during the entire course of parasite infection didn’t change parasitemia, mice weight or survival rate (Figures S5A-3C).

**Figure 5.**
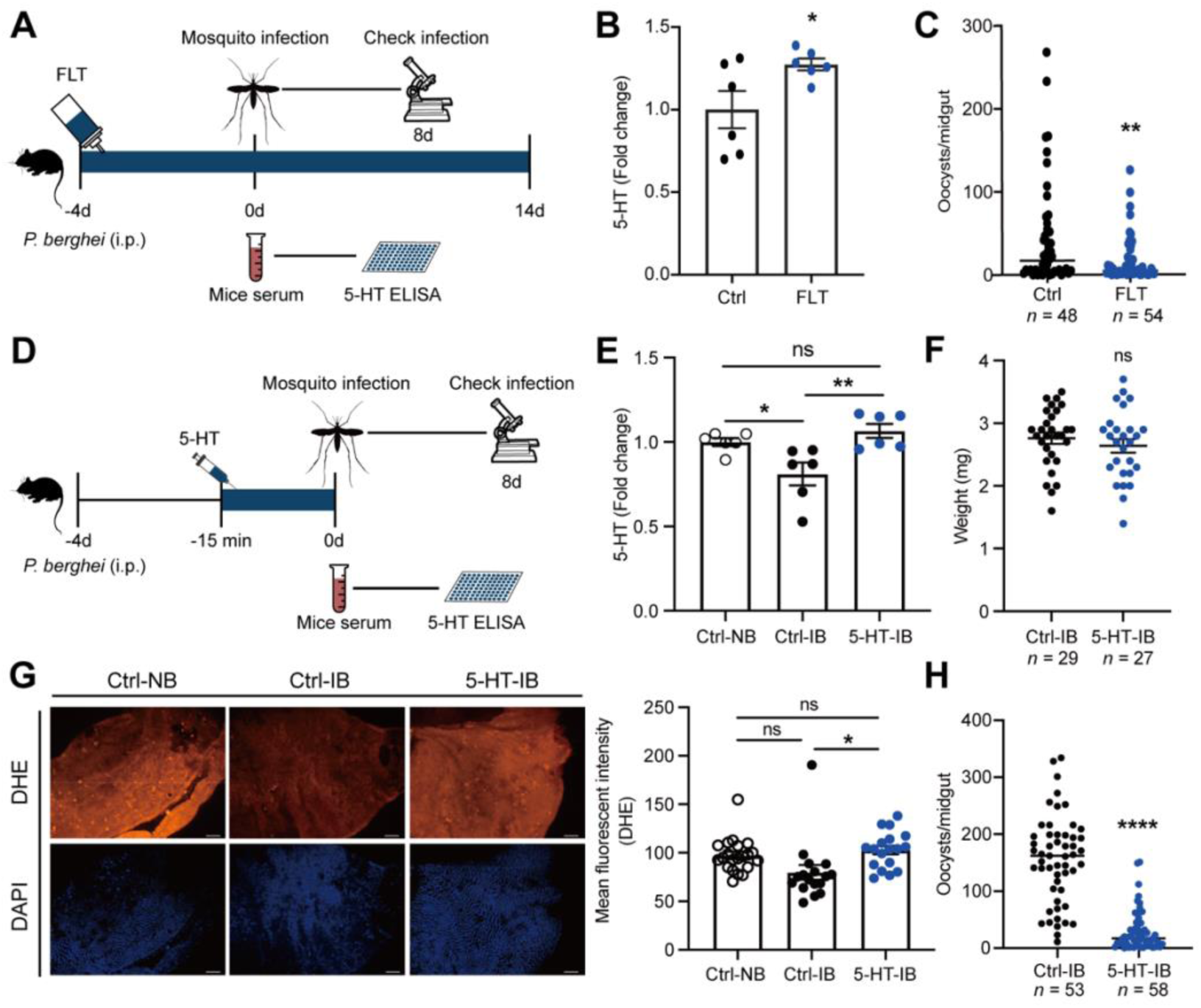
The influence of 5-HT in mice serum on *Plasmodium* infection in mosquitoes. (A) Schematic overview of fluoxetine supplementation in mice. Saline solution-treated mice were used as controls. (B) Fold change of 5-HT levels in the sera of control (*n* = 6) and fluoxetine (FLT, *n* = 6) treated mice 4 days post infection analyzed by ELISA. The 5-HT level in fluoxetine treated mice was normalized to that of controls. Each dot represented an individual mouse. Data were pooled from two independent experiments and shown as mean ± SEM. (C) Oocyst numbers in the midguts of mosquitoes fed on FLT treated (*n* = 54) or non-treated (*n* = 48) mice. Each dot represents an individual mosquito. Data were pooled from two independent experiments and horizontal lines represent the medians. (D) Schematic overview of 5-HT injection in mice. Saline solution-treated mice were used as controls. (E) Fold change of 5-HT levels in the sera of non-infected (Ctrl-NB, *n* = 6), *Plasmodium* infected (Ctrl-IB, *n* = 6) and *Plasmodium* infected mice treated with 5-HT (5-HT-IB, *n* = 6) 4 days post infection. The 5-HT levels were measured by ELISA. The 5-HT abundance in *Plasmodium* infected and 5-HT treated mice was normalized to that of controls. Each dot represented an individual mouse. Data were pooled from two independent experiments and shown as mean ± SEM. (F) The weight of fully engorged mosquitoes fed on *P. berghei*-infected mice injected with or without 5-HT. Each dot represented an individual mosquito. Data were pooled from two independent experiments and shown as mean ± SEM. (G) DHE (red) staining in the midguts of mosquitoes fed on non-infected, *Plasmodium* infected and *Plasmodium* infected + 5-HT treated mice 24 h post infection. Nuclei were stained with DAPI (blue). Representative images were shown (left). Mean fluorescence intensity was measured and calculated (right). Each dot represented an individual mosquito midgut. Data were pooled from two independent experiments and shown as mean ± SEM. Scale bar, 25 μm. (H) Oocyst numbers in the midguts of mosquitoes fed on mice injected without/with 5-HT. Each dot represents an individual mosquito. Data were pooled from two independent experiments and horizontal lines represented the medians. Significance was determined by two-sided Student’s t test in (B) and (F), Mann-Whitney test in (C) and (H) and ANOVA with Tukey’s test in (E) and (G). *p < 0.05, **p < 0.01, ****p < 0.0001, ns, not significant.

Fluoxetine has multiple influences on host physiology, including but not limited to modulating gut microbiota^31^ and immunity^32^. These effects might neutralize its inhibitory effect on *Plasmodium*. To verify the role of 5-HT in parasite infection in mosquitoes, we next injected 5-HT through tail vein of mice 15 mins before mosquito feeding (Figure 5D). 5-HT injection restored serum 5-HT levels to normal level (Figure 5E), but had no significant influence on mosquito blood ingestion (Figure 5F). As expected, mosquitoes that ingested elevated 5-HT showed increased levels of 5-HT and ROS in midguts compared to 5-HT non-ingested controls, and their 5-HT and ROS levels were similarly to mosquitoes that fed on normal blood (Figure 5G and S5D). The oocyst number was reduced from 162 in 5-HT non-ingested controls to 17 in 5-HT ingested ones (Figure 5H). Altogether, our data show that reversing the *Plasmodium*-mediated 5-HT reduction in host serum effectively suppresses *P. berghei* infection in mosquitoes.

## DISCUSSION

5-HT is a biogenic amine that plays a role in various physiological processes. In mammals, 5-HT produced within the central nervous system regulates mood, behavior, appetite and energy expenditure. The peripheral 5-HT, mainly generated by the gut, contributes to energy metabolism in multiple organs.^33^ In this study, we demonstrate that malaria parasite infection decreases the level of 5-HT in mammalian blood. The reduced 5-HT acquired from mice during blood feeding fails to efficiently elicit ROS generation in the mosquito midgut, thereby facilitating *Plasmodium* infection in *Anopheles* mosquitoes. Dietary interventions that increase 5-HT levels in mosquitoes through sugar and blood meals both suppress parasite infection in mosquitoes.

The role of peripheral 5-HT in *Plasmodium* pathogenesis in mammals remains unclear. Malaria parasites are known to suppress host immune responses by inducing the expression of indoleamine 2,3 dioxygenase (IDO), which is the rate-limiting enzyme of the kynurenine pathway in mammals.^34–36^ The metabolites along the kynurenine pathway have been implicated in the pathogenesis of murine and human cerebral malaria.^37^ The shift in tryptophan metabolism towards the kynurenine pathway may lead to the reduction of 5-HT production. The influence of 5-HT on the development of blood stage *Plasmodium* in vitro has been reported, but the results are controversial. For example, studies have shown that 5-HT promotes the formation of schizonts in the human malaria parasite *P. falciparum* by inducing Ca^2+^ mobilization.^38^ In contrast, when the intraerythrocytic stage of *P. falciparum* is treated with 5-HT receptor agonists, parasite growth is inhibited by blocking its surface membrane channel.^39^ As a regulator of the immune system, 5-HT also modulates the activation and function of multiple immune cells. However, its immune activating or suppressing effect is context dependent.^40^ Multidirectional interactions between 5-HT, mood and the peripheral immune system have been observed in viral and bacterial diseases,^40^ suggesting a potential link between 5-HT and *Plasmodium* pathogenesis.

5-HT is also an important neurotransmitter and neuromodulator in mosquitoes. It regulates hearing, heart rate, development, reproduction, metabolism, blood-feeding and flight behaviors of adult mosquitoes.^41–44^ Here we show that 5-HT also regulates the infection outcome of *P. berghei* in *A. stephensi* by modulating mitochondrial ROS production. In mammals, 5-HT is converted to ROS through the mitochondrial enzyme monoamine oxidase-A (MAO-A).^26,45^ It is possible that increasing the uptake of 5-HT in mosquitoes could also directly enhance ROS production. Additionally, we found that 5-HT accumulates dysfunctional mitochondria through inhibition of mitophagy, thereby aggravating ROS generation. Consistent with our findings, an increased uptake of 5-HT results in an elevated ROS production, leading to mitochondrial damage. This damage causes premature senescence and the pathogenesis of steatohepatitis in mammals.^46^ However, the mechanism by which 5-HT inhibits mitophagy remains unclear. In human hepatocellular cancer, 5-HT activates downstream signals, p70S6K and 4E-BP1, of the mammalian target of rapamycin (mTOR) in a mTOR-independent manner, and it inhibits autophagy.^47^ In mice cardiomyocytes, the activation of MAO-A leads to the accumulation of p53. This accumulation inhibits the translocation of parkin, a key factor that regulates mitophagy, from the cytoplasm to the mitochondria, ultimately leading to the inhibition of mitophagy.^26^ Further studies will be needed to investigate the mechanisms of 5-HT-mediated mitophagy inhibition.

Moreover, the mechanisms underlying the role of mitochondrial ROS in the elimination of *P. berghei* remain unclear. Mitochondrial ROS can trigger NADPH oxidase-mediated cellular ROS generation in mammalians and plants.^48^ Here we show that sequestration of mitochondrial ROS inhibits the total ROS generation, while blocking the NOX-derived ROS doesn’t influence mitochondrial ROS. These results indicate that mitochondrial ROS in mosquitoes may similarly play a role in promoting cellular ROS generation, which in turn influences the survival of *Plasmodium*.

Animal blood is crucial for mosquito physiology and reproduction, as it serves as their primary source of nutrition.^49^ Moreover, blood constituents have been increasingly recognized as important regulators for vector competence. For example, human low-density lipoprotein inhibits dengue virus acquisition in mosquitoes.^50^ Human blood-derived miRNA, hsa-miR-150-5p, disseminates to mosquito hemocoel and facilitates dengue virus infection by suppressing the expression of the antiviral *chymotrypsin* gene in mosquitoes.^51^ Our study reveals a novel role of a blood-derived metabolite, 5-HT, in modulating the vector competence of mosquitoes for parasite infection. Elevating the 5-HT level in mouse serum restores the 5-HT level in mosquitoes and increases their ability to eliminate parasites. In line with our findings, there is a negative correlation between serum iron levels in humans and dengue virus acquisition by mosquitoes. Elevating serum iron concentration in mice reduces dengue virus infection in *Aedes* mosquitoes.^52^ Interestingly, although increased uptake of 5-HT induces ROS generation and accumulates dysfunctional mitochondria, we didn’t find any defects in mosquito feeding capacity or survival. One possible explanation is that a moderate increase in ROS generation induced by 5-HT helps mosquitoes eliminate parasite infection without negatively affecting their physiology. Altogether, these findings suggest the potential for manipulating host metabolism to suppress pathogen transmission in vectors.

## STAR★METHODS

Ethics statement: This study was reviewed and approved by the Institutional Review Board of Shandong Institute of Parasitic Diseases, China. Informed consent was obtained from all participant. All blood samples were collected for the standard diagnostic tests, with no additional burden to the patients. All procedures involving mosquitoes and mice were carried out according to the guidelines for animal care and use of Fudan University and were permitted by the Animal Care and Use Committee, Fudan University, China.

Mosquito rearing and treatments: *A. stephensi* (strain Hor) was reared in the insectary with 28°C, 80% relative humidity and 12:12 light/dark cycles. Adults were fed on 10% sucrose solution and females were fed on mice for laying eggs. The chemicals, including 5-HT, a-Methyl-DL-tryptophan (AMTP, Sigma), spermidine (SPD, Sigma), H_2_O_2_ (Sangon, China), Vitamin C (Vc, Sigma), MitoTempo (Sigma), Dibenziodolium chloride (DPI, Sigma), were dissolved in sterile water, and carbonyl cyanide m-chlorophenyl hydrazone (CCCP, Yeason, China) was dissolved in DMSO. Newly-emerged mosquitoes were fed with 10% sucrose solution containing 100 μM AMTP,^53^ 100 μM spermidine^54^, 50 nM CCCP, and 5-HT and H_2_O_2_ with dedicated concentrations for four days prior to blood feeding, respectively. For ROS inhibition, antioxidants, including 3.3 mM Vitamin C,^55^ 50 μM MitoTempo^56^ and 50 μM DPI^24^ were administrated along with 5-HT through water during 24 h starvation prior to blood feeding. To administrate 5-HT to mosquitoes through blood meal, three to five days old adult mosquitoes were fed on mice that were orally supplemented with fluoxetine or intravenously injected with 5-HT.

Cell cultures: Cell line MSQ43 was grown in Schneider’s medium (Gibco) supplemented with 10% heat-inactivated fetal bovine serum (FBS, Gibco), 100 IU/mL penicillin and 100 μg/mL streptomycin (Thermo Fisher) at 28 °C. For 5-HT treatment, approximately 5 × 10^5^ cells were seeded per well in 12-well plates and incubated with 1 μM 5-HT for 3 days until they reached 70–90% confluency, and then used for subsequent detection. The mitophagy inducer CCCP was used as a positive control as described.^57^

*P. berghei* infection: Six to eight-week-old Balb/c mice were injected intraperitoneally (i.p.) with 10^6^ infected RBCs with *P. berghei* (ANKA).^58^ To evaluate parasitemia, thin blood smears were taken for Giemsa staining (Baso Diagnostics Inc, Zhuhai, China) daily from day 3 post injection. When the parasitemia reached 4-6%, the infected mice were used for mosquito infection. The engorged mosquitoes were maintained at 21°C. The unengorged mosquitoes were removed 24 h post blood meal. To evaluate the infection status, mosquito midguts were dissected at the indicated time points after infection. At 15 min post-infection, the gamete levels were determined using qPCR. At 12 h post-infection, the retort numbers were counted by examining thin blood smears from midguts containing the blood bolus. At 24 h post-infection, the blood bolus was removed from the midguts. After multiple PBS washes, the ookinete number in the midgut epithelium was examined using a fluorescence microscope. At 8 days post-infection, oocyst numbers were counted microscopically.

Mice treatments: For fluoxetine treatment, fluoxetine (15 μg/ml) (Sigma) was dissolved in sterile water. Six to eight-week-old Balb/c mice right after intraperitoneally injected with *P. berghei* were given fluoxetine (15 μg/ml)-containing drinking water for 4 days and 14 days, respectively. Control mice with provided with the saline solution as described.^59^ Parasitemia, 5-HT level, and weight of these mice were examined at dedicated time. For administration of 5-HT intravenously, 5-HT stock solution (1 mM) was prepared in sterile water and diluted to a final working concentration in saline solution. Mice with 4-6% parasitemia were injected with 5-HT via the tail vein at 0.5 mg/kg. Mice injected with equal amount of saline solution were used as control.^60^ Mosquitoes were allowed to feed 15 min post 5-HT administration.

5-HT measurement: The 5-HT levels of mosquitoes were measured using a serotonin ELISA kit (Biovision, USA) according to the manufacturer’s instructions. In brief, 30 midguts with blood bolus and 60 midguts without blood bolus, which were removed 24 h post blood meal were pooled for one biological sample. 30 midguts 3 days post blood meal were pooled for one biological sample. 25 whole mosquitos 4 days post treatment (24 h prior to blood meal) were pooled for one biological sample. Each sample was homogenized in 450 μL PBS and stored at −20°C overnight. Two freeze-thaw cycles were performed to break the cell membranes, and the homogenates were centrifuged for 5 min at 5000 × g. The supernatant was used for 5-HT quantification immediately.

LC-MS analysis: The blood samples from both normal and *Plasmodium*-infected mice and humans were collected in 1.5 mL EP tubes and allowed to settle for at least 1 h at 37 °C. Afterward, the blood samples were kept at 4 °C overnight to ensure complete blood clotting. The serum was then separated by centrifugation at 2000 x g for 10 min. The serum samples were sent to APExBIO Technology LLC in China for liquid chromatography-mass spectrometry (LC– MS) analysis using a Nexera UHPLC system (Shimadzu) coupled to an ABSciex QTrap 5500 or 6500 mass spectrometer (Framingham). Peak identification and amounts of metabolites were evaluated using Analyst and SCIEX OS software based on the known amounts of tryptophan metabolites.

Hemoglobin quantification: The hemoglobin levels of mosquitoes were measured using a hemoglobin assay kit (Abcam) according to the manufacturer’s instructions. In brief, 30 midguts with blood bolus 0 h post blood meal were pooled for one biological sample. Each sample was homogenized in 500 μL distilled water and used for hemoglobin quantification immediately.

ROS detection: Superoxide anion levels were detected in live tissues as previously described.^61^ In brief, midguts 15min and 24 h post blood meal were dissected in PBS and stained with 5 μM of the intracellular ROS-sensitive dye Dihydroethidium (DHE, Beyotime, China) for 20 min at room temperature in dim light, followed by 3× washings in PBS for 15 min. Then the midguts were stained with 4’,6’-diamidino-2-phenylindole (DAPI) (Solarbio, China) for 10 min and mounted using FluoromountTM Aqueous Mounting Medium (Sigma-Aldrich, USA). Images were acquired using a fluorescence microscope (Olympus, Germany). The same exposure parameters were used to compare fluorescence levels in different samples. Mean fluorescence intensity from the whole midgut was measured and calculated by Image J.

For hydrogen peroxide (H_2_O_2_) measurement, midguts 24 h post infection were dissected and assessed by H_2_O_2_ detection Kit (Beyotime, China) according to the protocol. Briefly, 15 midguts were pooled for one biological replicate and homogenized in lysis buffer provided by the kit. A hundred μl supernatant of the homogenate after centrifugation was measured at OD560 nm using a multiwell plate reader (Synergy™ 2, BioTek). The midguts protein levels were determined by BCA assay (Thermo Fisher). The H_2_O_2_ levels were normalized to protein amount.

To measure the mitochondrial ROS level, midguts 15 min and 24 h post infection were dissected in PBS and incubated with 5 μM of the mitochondrial superoxide Indicator, MitoSox Red (Yeasen, China) for 30 min at 37 °C in dim light, followed by 3× washings in PBS for 15 min. Then the midguts were stained with DAPI for 10 min and mounted using Mounting Medium. Images were acquired using a fluorescence microscope (Olympus, Germany). The same exposure parameters were used to compare fluorescence levels in different samples. Mean fluorescence intensity from the whole midgut was measured and calculated by Image J.

Mitochondrial membrane potential measurement: To assess mitochondrial membrane potential, midguts were dissected 24 h post infection in PBS and incubated in 100 nM TMRM dye (Invitrogen) for 30 min at 37 °C in dim light, followed by 3× washings in PBS for 15 min. Then the midguts were stained with DAPI for 10 min and mounted using Mounting Medium. Images were acquired using a fluorescence microscope (Olympus, Germany). The same exposure parameters were used to compare fluorescence levels in different samples. Mean fluorescence intensity from the whole midgut was measured and calculated by Image J.

Oxygen consumption rate measurement: Mitochondrial respiration of MSQ43 cells were monitored at 25 °C using the Oxygraph-2k (Oroboros) according to the operating instruction.^62^ In brief, 1×10^5^ cells cultured in 6-well plate were collected and resuspended in 200 μl serum-free Schneider’s medium (Gibco). Cells were equilibrated for 20 minutes in 2.5 ml medium prior to measurements. For analyzing the respiration of each mitochondrial complex, the following compounds were then sequentially injected to the chamber: 0.25mM oligomycin, 0.1mM FCCP, 0.5 μM rotenone and 2.5 μM antimycin A.^63^ The oxygen consumption was expressed as pmol O_2_ consumed per minutes per mg protein cells. The protein levels were determined using BCA assay (Thermo Fisher).

Transmission electron microscopy: Midguts of mosquitoes supplemented with 5-HT for four days were dissected at day four (24 hr prior to) and day 6 (24 hr post infection) in cool PBS and prefixed with 2.5% glutaraldehyde (Sangon, China) at 4°C overnight. After 3× washings in PBS for 15 min, midguts were post-fixed in 1 % osmium tetroxide (Sigma) for 2 h at 4°C, followed by dehydrating in an ascending series of ethanol (50%, 70%, 80%, 90%, and 100%). After dehydration, the samples were embedded in Epon 812 resin (EMCN, China) and polymerized at 65°C for 48 h.^64^ After trimming, blocks were sectioned in an Ultracut Reicher ultramicrotome. Regions of interest were selected, cut into ultrathin sections (50-nm thick) mounted on the copper grids, and then stained with uranyl acetate and lead citrate. The sections were examined and photographed in a Jeol JEM 1400 electron microscope performed by Servicebio Technology LLC, China. To quantify mitochondrial number, size and the percentage of mitochondria with abnormal cristae, scanned images of at least 3 sections of each midgut cell were analyzed using Fiji ImageJ (NIH).^65^

Citrate synthase activity assay: The activity of citrate synthase was measured as described.^66^ In brief, 100 midguts of sugar fed mosquitoes administrated with/without serotonin were dissected in cool PBS and homogenized in 200 μl lysis buffer (0.25% TritonX-100/PBS) at 4°C. After 1: 2 dilution in lysis buffer, 40 μl of the lysate mixed with 60 μl reaction buffer (0.25% TritonX-100/PBS, 0.31mM acetyl CoA, 0.1mM DTNB and 0.5mM oxaloacetate). The activity of citrate synthase was measured at 412 nm on a regular kinetic program (every 30 s for 5 min) at 30°C immediately by a multiwell plate reader (Synergy™ 2, BioTek).

Lysotracker staining: Midguts were dissected 24 h post infection in PBS and incubated in 1 μM Mito-Tracker Green (Beyotime) for 30 min at 37 °C in dim light, then the midguts were stained with 1 μM Lysotracker DS Red DND-99 (Invitrogen) for 5 min at room temperature in dim light, followed by 3× washings in PBS for 15 min. Then the midguts were stained with 4’,6’-diamidino-2-phenylindole (DAPI) (Solarbio, China) for 10 min and mounted using Fluoromount^TM^ Aqueous Mounting Medium (Sigma-Aldrich, USA). Images were acquired using a fluorescence microscope (Olympus, Germany). The same exposure parameters were used to compare fluorescence levels in different samples. Mean fluorescence intensity from the whole midgut was measured and calculated by Image J.

RNA interference: The cDNA clones of PINK1(ASTE000869) and plasmid eGFP (BD Biosciences) were served as templates for double-stranded RNA (dsRNA) preparation using gene-specific primers (Table S2). The dsRNA was synthesized by MEGAscript^TM^ T7 Transcription Kit (Thermo Fisher). Four to six-day-old females were injected intrathoracically with 69 nl of 4 μg/μl dsPINK1 using a Nanoject II microinjector (Drummond). Equal amounts of dsGFP were injected as a control. Silencing efficiency was examined two days post-dsRNA treatment by quantitative PCR as described below.

Quantitative PCR: For gene expression analysis in *A. stephensi*, total RNA was extracted from mosquitoes 15 min and 24 h post infection by TRIzol (Accurate Biology, China). Reverse transcription and quantitative PCR were performed as previously described.^58^ The expression levels of target genes were normalized by the *A. stephensi* ribosomal gene *S7*. The primers used for this study are listed in Table S2, Supporting Information.

Western blot: Proteins of 10 mosquitoes 24 h post infection were extracted in 300 μl lysis buffer (125 mM Tris, pH 6.8; 8 M urea; 2% SDS; 5% beta mercaptoethanol). Immunoblotting was performed using standard procedures using mouse anti-TOMM20 (Santa Cruz) (1:100), rabbit anti-LC3B (1:1000) (Abmart, China), and rabbit anti-actin (1:1000) (Abbkine, China). Intensity of the signals was quantified by Image J.

Immunohistochemistry: MSQ43 cells were fixed in 4% paraformaldehyde for 2 h at 4 °C, followed by three 10-min washes in PBS containing 0.1% Trixon-100. After blocking in 3% BSA for 2 h at 4 °C, cells were incubated with anti-TOMM20 mouse polyclonal antibody (Santa Cruz) (1:100 dilution) and anti-LC3B rabbit polyclonal antibody (Abcam) (1:50 dilution) overnight at 4 °C. The secondary antibody, anti-rabbit Alexa Fluor 546 and anti-mouse Alexa FITC 488 (Invitrogen) were used at 1:1000 dilution. The nucleus was stained with 10 μg/μl DAPI. Images were acquired by a Zeiss-LSM880 confocal microscope with Airyscan. The same exposure parameters were used to compare fluorescence levels in different images. The Pearson’s coefficient indexes between LC3 II and TOMM20 fluorescence intensities, the number of LC3 II puncta and the percent of LC3 II puncta with mitochondria were measured and calculated by Image J, respectively.

Statistical analysis: Replicates and sample sizes for all experiments were provided in the corresponding figure legends. All statistical analyses were performed using GraphPad Prism software (v.8). The comparison of two groups were analyzed using the Mann-Whitney test for non-normally distributed data, and Student’s t-test for normally distributed data. A Log-rank (Mantel-Cox) test was performed to compare the survival curves of *A. stephensi* exposed to 5-HT, H_2_O_2_ and control solution and mice supplemented with or without fluoxetine. The one-way ANOVA with different multiple comparisons tests were used to compare the difference among more than two groups depending on the normality of the data.

## SUPPLEMENTAL INFORMATION

Supplemental information can be found online.

## ACKNOWLEDGMENTS

This work was supported by National Natural Science Foundation of China (U1902211), the Shanghai Pilot Program for Basic Research - Fudan University (22TQ015) to J. W.

## AUTHOR CONTRIBUTIONS

Conceptualization, L.G., B.G., and J.W., Methodology, L.G., B.G., Y.B., W.X., S.B. and J.W.; Investigation, L.G., B.G., Y.B., W.X., S.B. and J.W.; Formal Analysis, L.G., and J.W., Writing Original Draft, L.G., and J.W.; Writing, Review & Editing, L.G., J.W.; Visualization, L.G., and J.W.; Funding Acquisition, J.W.; Resources, J.W., Supervision, J.W.

## DECLARATION OF INTERESTS

The authors declare no competing interests.

### Data Availability Statement

The data that support the findings of this study are available from the corresponding author upon reasonable request.

## Notes

### Competing Interest Statement

The authors have declared no competing interest.

