## supplemental figures for "Host 5-HT affects *Plasmodium* transmission in mosquitoes via modulating mosquito mitochondrial homeostasis"

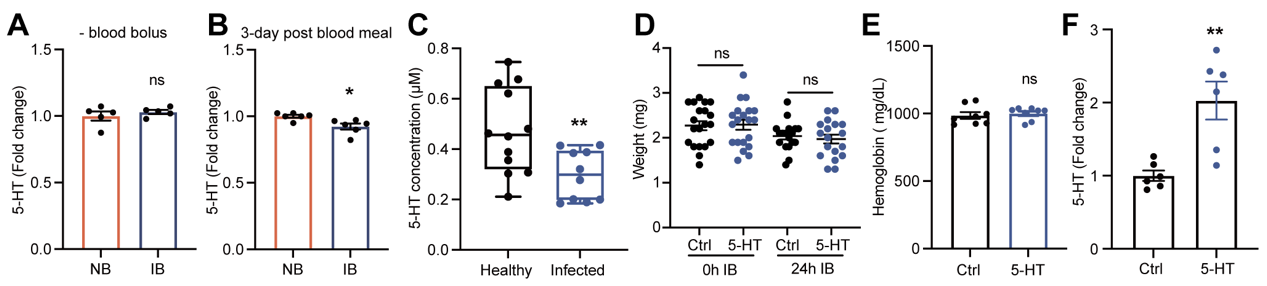

**Figure S1. The influence of 5-HT supplement on mosquito feeding capacity.**

(A) Fold change of 5-HT levels in the mosquito midguts 24 h post normal blood (NB, *n* = 5) and *P. berghei* containing infectious blood (IB, *n* = 5) analyzed by ELISA. The blood bolus was removed from the midgut 24h post blood meal. Sixty midguts were pooled for one sample. Each dot represented one biological replicate. Data were pooled from two independent experiments and shown as mean ± SEM.

(B) Fold change of 5-HT levels in the mosquito midguts 3 days post normal blood (NB, *n* = 6) and *P. berghei* containing infectious blood (IB, *n* = 6) analyzed by ELISA. Results represented that the blood was digested completely in the midgut 3 days post blood meal. Thirty midguts were pooled for one sample. Each dot represented one biological replicate. Data were pooled from two independent experiments and shown as mean ± SEM.

(C) 5-HT concentrations in the sera of healthy (Healthy, *n* = 12) and *Plasmodium* infected adults (Infected, *n* = 10). Each dot represented an individual and the data were shown as mean ± SEM.

(D) The weight of control and 1 μM 5-HT treated mosquitoes at 0 h (Ctrl, *n* = 20, 5-HT, *n* = 20) and 24 h (Ctrl, *n* = 18, 5-HT, *n* = 18) post infection. Each dot represented an individual mosquito. Data were pooled from two independent experiments and shown as mean ± SEM.

(E) Hemoglobin concentrations in the control (Ctrl, *n* = 8) and 1 μM 5-HT (5-HT, *n* = 8) treated mosquitoes (*n* = 6) at 0 h post infection. Each dot represented 30 mosquito midguts and the data were shown as mean ± SEM.

(F) Fold change of 5-HT levels in the control (*n* = 6) and 1 μM 5-HT treated mosquitoes (*n* = 6) 4 days post 5-HT treatment (24 h prior to blood feeding). The 5-HT abundance in 5-HT treated mosquitoes was normalized to that of controls. Twenty-five mosquitoes were pooled for one sample. Each dot represented one biological replicate. Data were pooled from two independent biological experiments and shown as mean ± SEM.

Significance was determined by two-sided Student’s t test. *p < 0.05, **p < 0.01, ns, not significant.

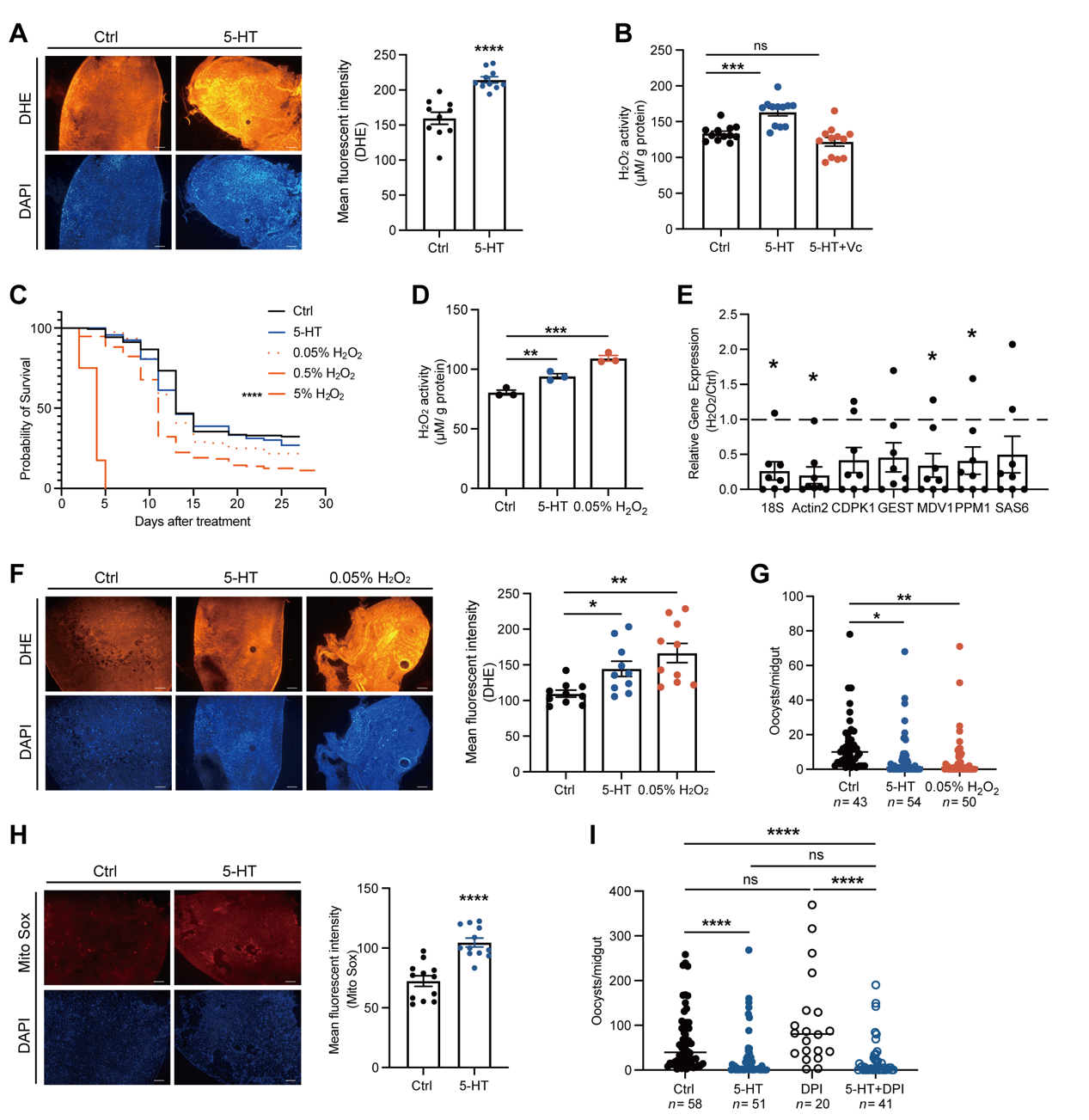

**Figure S2. The influence of H_2_O_2_ on *P. berghei* infection in *A. stephensi.***

(A) DHE staining (red) in the midguts of control and 5-HT (1 μM) treated mosquitoes 15 min post infection. Nuclei were stained with DAPI (blue). Representative images were shown (left). Mean fluorescent intensity was measured and calculated (right). Each dot represented an individual mosquito midgut. Data were pooled from two independent experiments and shown as mean ± SEM. Scale bar, 25 μm.

(B) The levels of H_2_O_2_ in the midguts of control, 5-HT and 5-HT + Vc treated mosquitoes 24 h post infection. Each dot represented an individual mosquito. Data were pooled from three independent experiments and shown as mean ± SEM.

(C) Survival assay of mosquitoes treated with 5-HT and different concentrations of H_2_O_2_ (n = 40–159 mosquitoes per group). Results were pooled from two independent experiments.

(D) The levels of H_2_O_2_ in the midguts of control, 5-HT- (1 μM) and 0.05% H_2_O_2_- treated mosquitoes 24 h post infection. Each dot represented an individual mosquito. Data were pooled from three independent experiments and shown as mean ± SEM.

(E) Fold changes of male gametogenesis associated genes in the midguts of control (*n* = 8) and 0.05% H_2_O_2_- (*n* = 8) treated mosquitoes 15 min post infection. The expression level of the target gene was normalized to *S7*. The relative expression level of target genes in 0.05% H_2_O_2_- treated mosquitoes was normalized to that in controls. Each dot represented five mosquito midguts. Data were shown as mean ± SEM.

(F) DHE staining (red) in the midguts of control, 5-HT- (1 μM) and 0.05% H_2_O_2_- treated mosquitoes 15 min post infection. Nuclei were stained with DAPI (blue). Representative images were shown (left). Mean fluorescent intensity was measured and calculated (right). Each dot represented an individual mosquito midgut. Data were pooled from two independent experiments and shown as mean ± SEM. Scale bar, 25 μm.

(G) Oocyst numbers in the midguts of Control (*n* = 43), 5-HT- (*n* = 54) and 0.05% H_2_O_2-_ (*n* = 50) treated mosquitoes. Each dot represents an individual mosquito. Data were pooled from two independent experiments and horizontal lines represented the medians.

(H) MitoSOX (red) staining in the midgut of control and 5-HT treated mosquitoes 15 min post infection. Nuclei were stained with DAPI (blue). Representative images were shown (left). Mean fluorescent intensity was measured and calculated (right). Each dot represented an individual mosquito midgut. Data were pooled from two independent experiments and shown as mean ± SEM. Scale bar, 25 μm.

(I) Oocyst numbers in the midguts of Control (*n* = 58), 5-HT (*n* = 51), DPI (*n* = 20) and 5-HT+DPI (*n* = 41) treated mosquitoes. Each dot represents an individual mosquito. Data were pooled from two independent experiments and horizontal lines represented the medians.

Significance was determined by two-sided Student’s t test in (A), (E) and (H), ANOVA with Dunn’s test in (B), (D), (F) and (G) and Tukey’s test in (I), and A Log-rank (Mantel-Cox) test in (C). *p < 0.05, **p < 0.01, ***p < 0.001, ****p < 0.0001, ns, not significant.

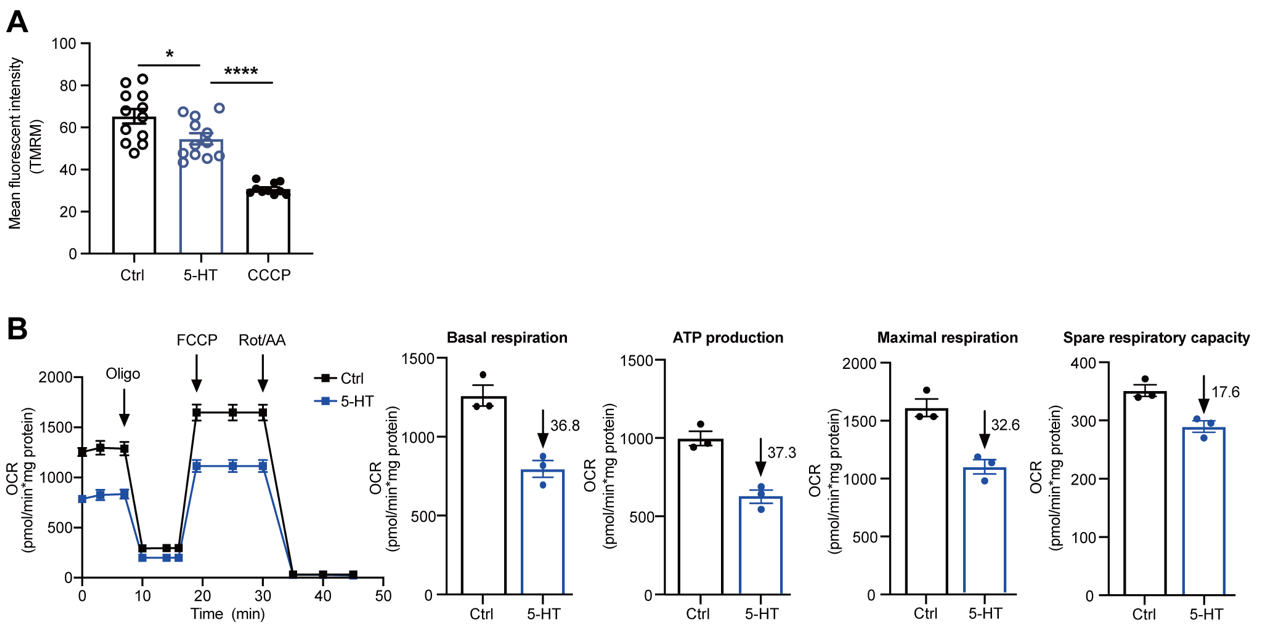

**Figure S3. The influence of 5-HT on MSQ43 cells.**

(A) The quantification of mitochondrial membrane potential of MSQ43 cells treated with 5-HT and CCCP were analyzed by TMRM staining. Each dot represented an independent experiment and the data were shown as mean ± SEM.

(B) Oxygen consumption rate (OCR) in control and 5-HT treated MSQ43 cells. Results were pooled from three independent experiments. Each dot represented an independent experiment and data were shown as mean ± SEM.

Significance was determined by ANOVA with Tukey’s test in (A) and two-sided Student’s t test in (B). *p < 0.05, ****p < 0.0001.

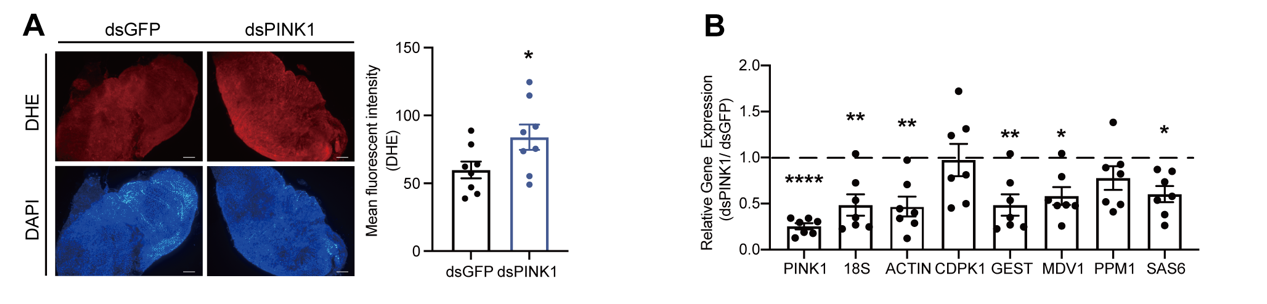

**Figure S4. The influence of *PINK1* on *P. berghei* infection in *A. stephensi.***

(A) DHE staining (red) in the midguts of dsGFP and dsPINK1 treated mosquitoes 15 min post infection. Nuclei were stained with DAPI (blue). Representative images were shown (left). Mean fluorescent intensity was measured and calculated (right). Each dot represented an individual mosquito midgut. Data were pooled from three independent experiments and shown as mean ± SEM. Scale bar, 25 μm.

(B) Fold changes of male gametogenesis associated genes in the midguts of dsGFP (*n* = 8) and dsPINK1 (*n* = 8) treated mosquitoes 15 min post infection. The expression level of the target gene was normalized to *S7*. The relative expression levels of target genes in dsPINK1 treated mosquitoes were normalized to those in controls. Each dot represented five mosquito midguts. Data were shown as mean ± SEM.

Significance was determined by two-sided Student’s t test in (A) and (B). *p < 0.05, **p < 0.01, ****p < 0.0001.

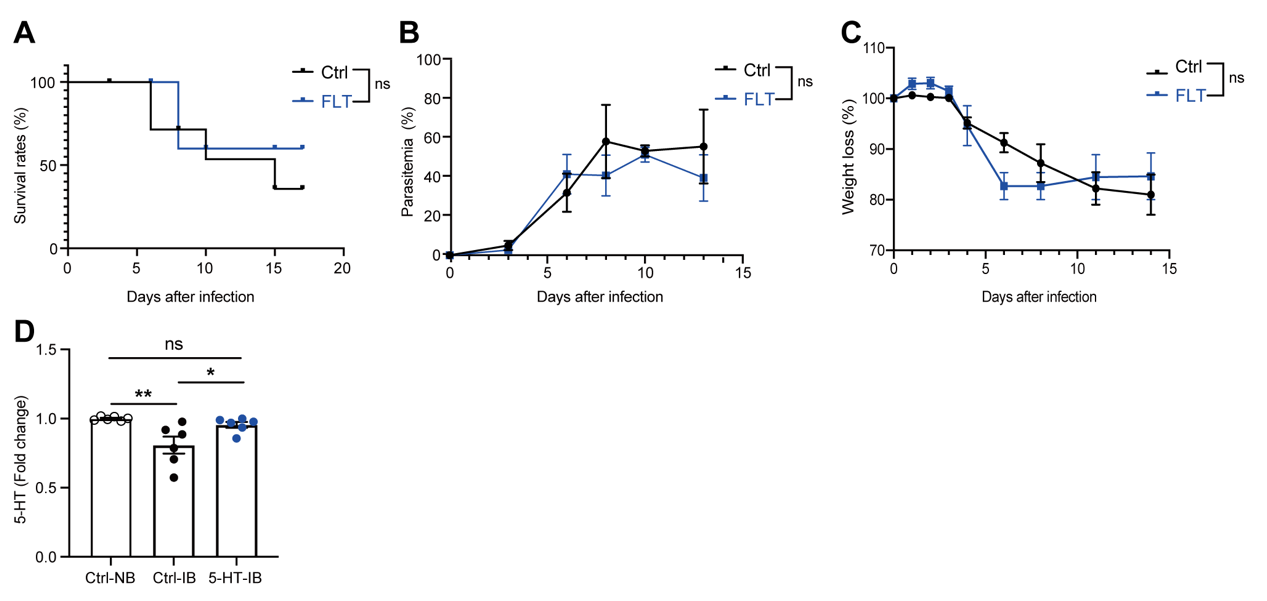

**Figure S5. The influence of fluoxetine treatment on *P. berghei* pathogenesis.**

(A-C) Survival (A , *n* = 8 mice per group), Parasitemia (B, *n* = 5 mice per group) and Weight loss (C, *n* = 5 mice per group) of mice following saline and fluoxetine treatment.

(D) Fold change of 5-HT levels in the midguts of mosquitoes fed on non-infected (Ctrl-NB, *n* = 6), *Plasmodium* infected (Ctrl-IB, *n* = 6) and *Plasmodium* infected + 5-HT injected (5-HT-IB, *n* = 6) mice 24 h post infection. The 5-HT abundance in *Plasmodium* infected and 5-HT treated mice was normalized to that of controls. Thirty midguts were pooled for one biological sample. Each dot represented one biological replicate. Data were pooled from two independent experiments and shown as mean ± SEM.

Significance was determined by A Log-rank (Mantel-Cox) test in (A), two-sided Student’s t test in (B) and (C), and ANOVA with Tukey’s test in (D). *p < 0.05, **p < 0.01, ns, not significant.
