## supplemental table 1 for "Host 5-HT affects *Plasmodium* transmission in mosquitoes via modulating mosquito mitochondrial homeostasis"

**Supplemental Table 1** The information of participants in this study

| **Subject ID** | **Sampling time** | **Sample ID** | **Age** | **Gender** | **Infected species** |
| --- | --- | --- | --- | --- | --- |
| Patient 1 | 2019.05.14 | I1 | 47 | Male | *P. malariae* |
| Patient 2 | 2019.05.15 | I2 | 34 | Male | *P. ovale* |
| Patient 3 | 2019.07.31 | I3 | 32 | Male | *P. falciparum* |
| Patient 4 | 2019.08.02 | I4 | 51 | Male | *P. falciparum* |
| Patient 5 | 2019.08.23 | I5 | 54 | Male | *P. ovale* |
| Patient 6 | 2019.09.20 | I6 | 29 | Male | *P. falciparum* |
| Patient 7 | 2020.12.24 | I7 | 54 | Male | *P. vivax* |
| Patient 8 | 2021.07.16 | I8 | 39 | Male | *P. ovale* |
| Patient 9 | 2021.07.01 | I9 | 35 | Male | *P. falciparum* |
| Patient 10 | 2021.06.24 | I10 | 53 | Male | *P. vivax* |
| Healthy 1 | 2020.12.10 | N3 | 60 | Male | / |
| Healthy 2 | 2020.12.15 | N4 | 39 | Male | / |
| Healthy 3 | 2020.12.15 | N6 | 61 | Male | / |
| Healthy 4 | 2020.12.17 | N8 | 44 | Male | / |
| Healthy 5 | 2020.12.17 | N9 | 32 | Male | / |
| Healthy 6 | 2021.09.01 | N15 | 27 | Male | / |
| Healthy 7 | 2021.09.01 | N18 | 26 | Male | / |
| Healthy 8 | 2021.09.01 | N19 | 26 | Male | / |
| Healthy 9 | 2021.09.01 | N20 | 44 | Male | / |
| Healthy 10 | 2021.09.01 | N21 | 27 | Male | / |
| Healthy 11 | 2021.09.01 | N22 | 26 | Male | / |
| Healthy 12 | 2021.09.01 | N23 | 26 | Male | / |
