## supplemental table 2 for "Host 5-HT affects *Plasmodium* transmission in mosquitoes via modulating mosquito mitochondrial homeostasis"

**Supplemental Table 2** qPCR primers used in this study

| Primer name  (accession #) | Primer sequences  (F, Forward; R, reverse) | Amplicon size  (bp) | | Tm  (˚C) | |
| --- | --- | --- | --- | --- | --- |
| ^∆^T7-GFP | F: 5' TAATACGACTCACTATAGGGTCAGTGGAGAGGGTGAAG 3' | | 454 | | 60 |
| (BD Biosciences) | R: 5' TAATACGACTCACTATAGGCTAGTTGAACGGATCCATC 3' | |  | |  |
| *PINK1  (ASTE00869) | F: 5' GAAGGATGGACAGAAGAAG3' | | 495 | | 60 |
|  | R: 5' CGTGGCAGACATTGATAA 3' | |  | |  |
| ^∆^T7-PINK1 | F: 5' TAATACGACTCACTATAGGGAAGGATGGACAGAAGAAG 3' | | 495 | | 60 |
| (ASTE00869) | R: 5' TAATACGACTCACTATAGGCGTGGCAGACATTGATAA 3' | |  | |  |
| qPINK1  （ASTE00869） | F: 5' TCGGTGAGAAGGATATTGAA 3' | | 168 | | 60 |
|  | R: 5' TATGGATGTTGTCGCTGTA 3' | |  | |  |
| qS7  （ASTE004816） | F: 5' TGCGGAGCGTCGTATTCTGC 3' | | 79 | | 60 |
|  | R: 5' ACACAGCGGTGAGCGTTCG 3' | |  | |  |
| qATT | F: 5' CGCCTCACCATTGTCAAGCC 3' | | 125 | | 58 |
| （ASTE009529） | R: 5' GTCCGTTCCGTATCCGTCCT 3' | |  | |  |
| qCEC  （ASTE007106） | F: 5' GCTGCTCTTTCTCGTTGCG 3' | | 98 | | 60 |
|  | R: 5' CGGCACCTTCCACCTTCT 3' | |  | |  |
| qGAM  （ASTE002252） | F: 5' CCGCTGTTCGTCCTCGTTCA 3' | | 91 | | 60 |
|  | R: 5' GCACACGGACGCCAACTTCT 3' | |  | |  |
| qDEF  （ASTE011281） | F: 5' CCGCCTTGAACACGCTCCT 3' | | 116 | | 60 |
|  | R: 5' GCTGCCGACACCGAATCCA 3' | |  | |  |
| qTEP1  （ASTE010227） | F: 5' CCTGGGTGCGTGGGAAAC 3' | | 106 | | 60 |
|  | R: 5' GCCTTGCTGTCGTTCGTGAT 3' | |  | |  |
| qPPO  （ASTE000587） | F: 5' TTCTGCGGTTGCGGCTGG 3' | | 106 | | 60 |
|  | R: 5' CGGCGTCTTGCTCGTAGTCG 3' | |  | |  |
| qNOS  （ASTE008593） | F: 5' CAGCGAACGGACGGCAAGCA 3' | | 186 | | 60 |
|  | R: 5' TGACACGACCAGCGGCAGGAT 3' | |  | |  |
| qDUOX  （ASTE003295） | F: 5' TCGTGAGCGTCGTCAGAAGC 3' | | 116 | | 60 |
|  | R: 5' CCTCACCGTCCAGCGATGC 3' | |  | |  |
| qPGRP-LA | F: 5' ACGCAGCCATCGGTGAGC 3' | | 131 | | 60 |
| (ASTE016413) | R: 5' GCAGACGGACAGTGTTCGGTTT 3' | |  | |  |
| qPGRP-LC | F:5' TGTGCCATCGTAGCGGTCAT 3' | | 98 | | 60 |
| （ASTE002618） | R:5' AGCCACTCGGTTCTCGTCAC 3' | |  | |  |
| qPGRP-LD | F: 5' GGCGTGTATCTGCTGCTGCT 3' | | 81 | | 60 |
| （ASTE010245） | R: 5' CCAGGCGGGTCGTTCCAC 3' | |  | |  |
| qCuSOD_3_ | F: 5' CACGAGTTCGGAGACAACACCAA 3' | | 143 | | 60 |
| （ASTE006711） | R: 5' ACCAAGTCCACCTTCGCTTCAC3' | |  | |  |
| qCAT | F: 5' TGGGATATGGTGGGCAACAATACG 3' | | 140 | | 60 |
| （ASTE010206） | R: 5' GCGGTCGGAGAACAGGAACATC 3' | |  | |  |
| qGSTO1 | F: 5' ACGGTTCAATGGTGCGGTCAT 3' | | 121 | | 60 |
| （ASTE003787） | R: 5' CGTGCCTTCAGCTCCTTCTCAA 3' | |  | |  |
| qUGT | F: 5' GACAGCCGAACGGTGGTGAAG 3' | | 121 | | 60 |
| （ASTE002225） | R: 5' CAGCAGCAGCAGCGACGAA 3' | |  | |  |
| qPbActin2 | F: 5' TTCTTGATAGTGGTGATGGCGTAA 3' | | 171 | | 60 |
| (PBANKA1030100) | R: 5' CGATTTCTCTTTCAGCGGTTGTAG 3' | |  | |  |
| qPbSAS6 | F: 5' AAAGGATATGGGCGTGATGGAT 3' | | 87 | | 60 |
| (PBANKA0106200) | R: 5' ACACTCATTAGATGTACCACCACT 3' | |  | |  |
| qPbGEST | F: 5' CGAATCAAATTAGTGAAGCATGGA 3' | | 146 | | 60 |
| (PBANKA1312700) | R: 5' TTCAGCGTCATTCTCAAGAAGA 3' | |  | |  |
| qPbCDPK1 | F: 5' TGGTGGGCAAAACGATCAAGA 3' | | 156 | | 60 |
| (PBANKA0314200) | R: 5' GCTTCCTCAGCCGTACATCT 3' | |  | |  |
| qPbPPM1 | F: 5' AGGGGATAGTCGCTGTGTCTT 3' | | 163 | | 60 |
| (PBANKA100700) | R: 5' ACCTCGGCATACTCCTAAGCAT 3' | |  | |  |
| qPbMDV1 | F: 5' TCCAACATCAACCATAGGGTGTCT 3' | | 112 | | 60 |
| (PBANKA1432200) | R: 5' TGCCTTGCCTCCACTTCCA 3' | |  | |  |
| qPb18S | F: 5' GGAGATTGGTTTTGACGTTTATGTG 3' | | 134 | | 60 |
| (PBANKA1245821) | R: 5' AAGCATTAAATAAAGCGAATA CATCCTTAC 3' | |  | |  |

*: Primers for gene cloning

^∆^: Primers for dsRNA synthesis

q: Primers for quantitative PCR
